# Dissemination of linezolid-resistance through sex pheromone plasmid transfer in *Enterococcs faecalis*

**DOI:** 10.1101/832691

**Authors:** Jiaqi Zou, Zhaobing Tang, Jia Yan, Hang Liu, Yingzhu Chen, Dawei Zhang, Jinxin Zhao, Yu Tang, Jing Zhang, Yun Xia

**Author notes:** Correspondence: Dr. Yun Xia, Department of Clinical Laboratory, the First Affiliated Hospital of Chongqing Medical University, 1 Youyi Rd, Chongqing 400016, China. Contributed equally to this work (Jiaqi Zou, Zhaobing Tang).

## Abstract

Despite recent recognition of the ATP-binding cassette protein OptrA as an important mediator of linezolid-resistance in *Enterococcus faecalis* worldwide, the mechanisms of *optrA* gene acquisition and transfer remain poorly understood. In this study, we performed comprehensive molecular and phenotypic profiling of 44 *optrA*-carrying *E. faecalis* clinical isolates with linezolid-resistance. Pulse-field gel electrophoresis and DNA hybridization revealed the presence of *optrA* in the plasmid in 26 (59%) isolates and in the chromosome in 18 (41%) isolates. Conjugation experiments showed a successful transfer of *optrA* in 88.5% (23/26) of isolates carrying *optrA* in plasmids while no transfer occurred in any isolates carrying *optrA* in the chromosome (0/18). All 23 transconjugants exhibited *in vitro* resistance to linezolid and several other antibiotics, and were confirmed to contain *optrA* and other resistance genes. Plasmid typing demonstrated a predominance (18/23 or 78%) of *rep*_9_–type plasmids (pCF10 prototype) known to be the best studied sex pheromone responsive plasmids. Full plasmid genome sequencing of one isolate revealed the presence of drug resistance genes (*optrA* and *fexA*) and multiple sex pheromone response genes in the same plasmid, which represents the first sex pheromone responsive plasmid carrying *optrA* from a clinical isolate. PCR-based genotyping revealed the presence of three key sex pheromone response genes (*prgA, prgB* and *prgC*) in almost all 23 *optrA*-carrying isolates tested. Finally, functional studies of these isolates by clumping induction assay detected different degrees of clumping in most of the 23 isolates. Our analysis strongly suggests that *optrA*-mediated linezolid-resistance can be widely disseminated through sex pheromone plasmid transfer.

## Introduction

*Enterococcus faecalis* is an opportunistic pathogen that resides primarily in the gastrointestinal tract of most healthy individuals, but can cause urinary tract infections, surgical infections and even fatal infections, such as endocarditis and bacteremia (1, 2). Its high-level intrinsic and acquired resistance to multiple drugs, especially vancomycin, has limited the treatment options (3, 4).

Linezolid, the first member of oxazolidinone antibiotic antibacterial agents, was approved for clinical use in 2000 as the last-resort drug for treatment of serious Gram-positive bacterial infections, including vancomycin-resistant *Enterococci* (VRE), methicillin-resistant *Staphylococcus aureus* (MRSA) and multi-drug resistant *Streptococcus pneumonia* (5). The increasing prevalence of linezolid-resistant *E. faecalis* presents a formidable challenge to clinical treatment (6, 7). The common mechanisms of linezolid resistance in *E. faecalis* include point mutations in the chromosomal 23S rRNA gene or protein-coding genes encoding the L3 and L4 ribosomal proteins (8, 9). Other newly identified mechanisms include the presence of plasmid-borne chloramphenicol-florfenicol resistance (*cfr*) gene (10) or ribosomal protection genes *optrA* (11) and *poxtA* (12).

*OptrA* encodes the ATP-binding cassette F (ABC-F) protein family and mediates resistance to phenicols and oxazolidinones through protection of the bacterial ribosome from antibiotic inhibition (13, 14). Following the first report of the *optrA* gene in 2015 from an *E. faecalis* isolate obtained from a blood sample of a Chinese patient (13), *optrA* has been found worldwide not only in *E. faecalis* and *E. faecium* (15–22) but also in other Gram-positive bacteria such as *Staphylococcus sciuri* (23, 24) and *Streptococcus suis* (25). In addition, *optrA* has been detected not only in various human clinical samples but also in diverse livestock samples such as cow milk, faeces or meat from pigs, chicken and ducks (26, 27) as well as environmental samples such as soils (28) and urban wastewater (29). Search of the PubMed database using *optrA* as the keyword in the abstract identified 2, 10, 15, 23, 26 papers from 2015 to this year (as of the date of submission of this manuscript), respectively, with the geographic origin of these reports expanded from only one country (China) in 2015 to 17 countries (from 5 continentals) this year. Such rapid, widespread dissemination of this resistance gene suggests a highly efficient dissemination capacity among animals, humans and the environment.

Indeed, there have been reports of easy transfer of *optrA*-carrying plasmids *in vitro* between *E. faecalis* and *E. faecium* (13) and between human- and pig-derived *E. faecalis* (30). Our previous transcriptomics (31) and proteomics (32) analyses of a linezolid-resistant *E. faecalis* strain P10748 consistently revealed a significant co-upregulation of *optrA* gene with several genes involved in mating and pheromone response, implying a localization of this gene in a sex pheromone plasmid allowing highly efficient resistance transfer. However, the location of the *optrA* gene and the sequences of the plasmid and chromosome of this strain were not determined.

The goals of this study were to assess the distribution and transferability of the *optrA* gene in a case series of *E. faecalis* clinical isolates, determine the whole plasmid genome sequence in the linezolid-resistant *E. faecalis* strain P10748, and identify mechanisms behind *optrA* transfer.

## Materials and Methods

### 2.1. Bacterial strains and antimicrobial susceptibility testing

A panel of 44 linezolid-resistant *E. faecalis* isolates was obtained from the First Affiliated Hospital of Chongqing Medical University, Chongqing, China (31, 33). All isolates showed minimum inhibitory concentrations (MICs) > 4 mg/L as determined by the broth microdilution method following the Clinical and Laboratory Standards Institute (CLSI) recommendations. The drug resistance phenotypes and multi-locus sequence types (MLST) of all these isolates have been determined in our previous study (33) (Table 1 and Supplemental Table 1). The *E. faecalis* ATCC 29212 isolate obtained from the American Type Culture Collection (Manassas, VA, USA) was served as a quality control strain. The *E. faecalis* JH2-2 isolate was kindly provided by Dr. Tieli Zhou (the First Affiliated Hospital of Wenzhou Medical University, China) and used as the recipient strain in conjugation experiment.

**Table 1.** Characteristics of linezolid-resistance plasmids in 26 *E. faecalis* isolates carrying *optrA*

| Isolate ID | Plasmid size (kb)<sup>a</sup> | Rep-family | Prototype | Transferred resistance gene | MLST<sup>a</sup> |
| --- | --- | --- | --- | --- | --- |
| P10748 | ~54 | 9 | pCF10 | *optrA, tetL, ermB, lnuB, fexA* | ST480 |
| EF-3015 | ~150 | 6, 9 | pS86, pCF10 | *optrA, tetL, ermB, lnuB fexA,* | ST828 |
| EF-7013 | ~60 | 9 | pCF10 | *optrA, tetL, ermB, lnuB, fexA,* | ST16 |
| EF-8194 | ~80 | 9 | pCF10 | *optrA, tetL ermB, lnuB fexA,* | ST16 |
| EF-9289 | ~80 | 1, 7, 9 | pIP501, pUSA02, pAD1 | *optrA, tetL, ermB, fexA* | ST826 |
| EF-3139 | ~100 | 1, 9 | pIP501, pCF10 | *optrA, tetL, tetM, ermB, lnuB fexA,* | ST386 |
| EF-5136 | ~55 | 9 | pCF10 | *optrA, tetL, tetM, ermB, lnuB fexA,* | ST16 |
| EF-2084 | ~80 | 1, 9 | pIP501, pCF10 | *optrA, tetL, tetM, ermB, lnuB, fexA,* | ST480 |
| EF-0132 | ~80 | 1, 9 | pIP501, pCF10 | *optrA, tetL ermB, lnuB, fexA,* | ST585 |
| EF-4003 | ~55, ~350<sup>b</sup> | 7 | pUSA02 | *optrA, tetL, tetM, ermB, fexA* | ST16 |
| EF-8042 | ~30, ~200<sup>b</sup> | 1, 9 | pIP501, pCF10 | *optrA, tetL, ermB, fexA* | ST69 |
| EF-0074 | ~65 | 7, 9 | pUSA02, pAD1 | *optrA, tetL, ermB, lnuB, fexA,* | ST826 |
| EF-6165 | ~80 | 1 | pIP501 | *optrA, tetL, tetM, ermB, lnuB, fexA,* | ST823 |
| EF-7094 | ~40, ~350<sup>b</sup> | 9 | pCF10 | *optrA, tetL, tetM, ermB, lnuB, fexA,* | ST632 |
| EF-1127 | ~55 | 1, 9 | pIP501, pCF10 | *optrA, tetL, tetM, ermB, lnuB, fexA,* | ST824 |
| EF-8137 | ~80 | 1, 7, 9 | pIP501, pUSA02, pCF10 | *optrA, tetL, tetM, ermB, lnuB, fexA,* | ST116 |
| EF-8282 | ~80 | 6,9 | pS86, pCF10 | *optrA, tetL, ermB, lnuB* | ST330 |
| EF-4245 | ~80 | 9 | pCF10 | *optrA, tetL, ermB, lnuB, fexA,* | ST823 |
| EF-1080 | ~40 | 9 | pCF10 | *optrA, tetL, tetM, ermB, lnuB, fexA* | ST69 |
| EF-2216 | ~80 | 1,9 | pIP501, pCF10 | *optrA, tetL, tetM, ermB, lnuB, fexA* | ST376 |
| EF-6166 | ~80 | NA | NA | *optrA, tetL, ermB, lnuB, fexA* | ST16 |
| EF-5015 | ~80 | NA | NA | *optrA, tetL, tetM, ermB, lnuB, fexA,* | ST480 |
| EF-2021 | ~90 | NA | NA | *optrA, tetL, ermB, lnuB fexA,* | ST714 |
| EF-8014 | ~55 | NT | NT | NT | ST632 |
| EF-3186 | ~30 | NT | NT | NT | ST69 |
| EF-1090 | ~55 | NT | NT | NT | ST585 |

<sup>a</sup>Multi-locus sequence types determined in our previous study (33).
<sup>b</sup>Two plasmids with different sizes.
NA, not available in plasmid typing with 19 known *rep* families. NT, not tested due to negative results in conjugation experiment.

### 2.2. Determination of the *optrA* location in plasmid or chromosome

To determine the location of *optrA*, S1 nuclease-pulsed field electrophoresis (S1-PFGE) and Southern blot were performed following the methods described previously (34, 35). Briefly, bacterial cells harvested from fresh culture were embedded in agarose gels and then digested by S1 nuclease (TaKaRa, Dalian, China) at 37 °C for 20 minutes. *Braenderup Salmonella* H9812 chromosomal DNA was digested with *XbaI* (TaKaRa, Dalian, China) for 4h at 37°C and used as the DNA size marker. Gel plugs were loaded onto 1% agarose gels and electrophoresed in the CHEF Mapper XA system (Bio-Rad, Hercules, CA) at 14℃, with a pulse time of 5–35 s, at 6 V cm-2 on a 120° angle in 0.5 × TBE for 20 h. Following transfer to Hybond N+ membranes (Amersham Biosciences, USA) by capillary blotting for more than 20 hours, blots were hybridized with digoxigenin (DIG)-labeled *optrA*-specific probe using the DIG High Prime DNA Labeling and Detection Starter Kit II (Roche Applied Sciences, Germany) following the manufacturer’s instructions.

### 2.3. Conjugation experiment and detection of resistance genes

To examine the transferability of plasmids containing *optrA*, filter mating was conducted on nitrocellulose filters (0.45um pore size, Millipore, USA) as described previously (36). Rifampicin-resistant *E. faecalis* JH2-2 was used as a recipient strain and *optrA*-positive *E. faecalis* was used as a donor strain. The donor and recipient strains were cultured to the exponential growth period (OD600 = 0.4-0.6) and then mixed at a ratio of 1:3. The mixture was shaken for 30 min and centrifuged. The pellet was placed on a filter for overnight incubation at 37℃. Transconjugants were selected on Brain Heart Infusion (BHI) agar (Solarbio, Beijing, China) supplemented with 25 mg/L of rifampin and 10mg/L of florfenicol. Subsequently, all transconjugants were subjected to PCR to confirm the presence or absence of the *optrA* gene.

To detect the transmission of resistance in transconjugants, antimicrobial susceptibility to linezolid, tetracycline, erythromycin, chloramphenicol and clindamycin of the transconjugants were determined as described above. Known potential antibiotic resistance genes, including *tetM*, *tetL*, *ermB, lnuB* and *fexA,* were screened by PCR amplification using the primers listed in Table S2 (37, 38). Positive PCR products were sequenced commercially by Sangon Biotech (Shanghai, China). Resulting sequences were analyzed using MegAlign (version 7.1.0, DNASTAR, USA) and compared with reference sequences in the NCBI Nucleotide Database.

### 2.4. Plasmid replicon typing

On the basis of plasmid classification schemes by Jensen et al (39), we designed PCR to detect 19 different types of *rep*-family plasmids (*rep*_1-11,_ *rep* _13-19_ and *rep* unique) in *E. faecalis* transconjugants. *Rep*_12_ family was not included owing to lacking a reference strain. All samples were first screened by multiplex PCR, followed by single-locus PCR for samples that were not clearly differentiated in multiplex PCR. *Rep*-families (*rep*_1_, *rep* _4_, *rep* _6-7_, *rep* _10_ and *rep*_13_) present in more than one species were defined as *rep*-families with a broad host range, whereas other *rep*-families (*rep*_2_*-*_3_, *rep* _5_, *rep* _8-9_, *rep*_11-12_ and *rep*_14-19_) present in only one species were defined as single-species specific *rep*-families. All positive PCR products were sequenced commercially and analyzed against sequences of known *rep*-families in the NCBI Nucleotide Database. In addition to transconjugants, all the *rep_-_*types were verified by PCR in original clinical isolates of *E. faecalis*.

### 2.5. Sequencing and analysis of the whole plasmid genome in P10748 harboring *optrA*

The whole plasmid genome in the *E. faecalis* strain P10748 was sequenced as a part of the *E. faecalis* genome sequencing project [U.S. National Center for Biotechnology Information (NCBI, Bethesda, MD, USA) BioProject accession no. SUB5737281]. Genomic DNA was extracted from overnight culture using the Qiagen Plasmid Midi Kit (Qiagen, Germany) according to the manufacturer’s instructions. The quantity and quality of genomic DNA were determined by NanoDrop spectrophotometer (Thermo Fisher Scientific) and agarose gel electrophoresis. Five micrograms of DNA were used to construct one Illumina library and sequenced using 250 base paired-end reads on the Illumina MiSeq platform commercially (Novogene Corporation, Hong Kong). A total of 20 million reads were obtained. After removal of low-quality reads and reads for *E. faecalis* nuclear genome sequences (GenBank accession no. CP008816), the remaining 285,862 reads were assembled using SeqMan Pro (version 14.1.0.118, DNASTAR, Madison, WI), resulting in a circular assembly of the complete plasmid genome. This assembly was validated by PCR and Sanger sequencing of multiple overlapping fragments, and by restriction mapping using *Nhe* I.

Gene prediction was conducted through a combination of multiple computational programs including GeneMarkS (40), RAST (41), ORF Finder (https://www.ncbi.nlm.nih.gov/orffinder/) and BLAST (http://blast.ncbi.nlm.nih.gov/Blast.cgi).

### 2.6. Distribution of key sex-pheromone-response genes in *optrA*-carrying *E. faecalis* isolates and their transconjugants

Genomic DNA was extracted from overnight cultures using the HiPure Bacterial DNA Kit (Magen, Guangzhou, China) according to the manufacturer’s protocol. The presence of three key sex-pheromone-response genes (*prgA, prgB* and *prgC*) in the donor and transconjugant strains was determined by PCR using primers listed in Table S2. PCR reaction was performed with the following thermocycling program: 95 °C for 5 minutes, 30 cycles of 95 °C for 1 minute, 50 °C for 30 seconds, 72 °C for 1 minute, and a final extension at 72 °C for 10 minutes. All positive PCR products were sequenced and the resulting sequences were blasted against the NCBI Nucleotide Database.

### 2.7. Determination of sex pheromone responses by clumping induction assay (CIA)

CIA was performed as described previously (42). Pheromone-containing filtrates were prepared from *E. faecalis* JH2-2 strain grown overnight in BHI broth (referred to as the recipient strain). The cells were pelleted by centrifugation, and the supernatant was filtered through a 0.22 μm filter membrane. The filtrates were autoclaved at 121 °C for 20 min, and stored at 4 °C before use. *E. faecalis* culture (from clinical isolates or their transconjugants, referred to as the donor strain) was added to 0.5 ml of pheromone-containing filtrate, followed by incubation at 37 °C for 3h with shaking. To test the type of sex pheromones involved in mating, synthetic sex pheromones cAD1 or cCF10 (Sangon Biotech, Shanghai, China) were used at the final concentration of 50 μg/ml instead of JH2-2-derived filtrates described above. Clumping status was monitored by visual inspection or microscope, and recorded by photography.

## RESULTS

### 3.1. Localization of *optrA* gene in the linezolid-resistant *E. faecalis*

Among 44 linezolid-resistant *E. faecalis* isolates analyzed by S1-PFGE, *optrA* was localized on the plasmid in 26 (59 %) isolates and on the chromosome in 18 (41%) isolates. The size of plasmids varied from 30 kb to 300 kb (Fig. 1). Of the 26 isolates with *optrA*-carrying plasmids, 23 had a single plasmid and the remaining 3 had a co-existence of two plasmids with different sizes (Table 1). The presence of *optrA* in all 44 isolates was confirmed by PCR using *optrA-*specific primers.

**Figure 1.**
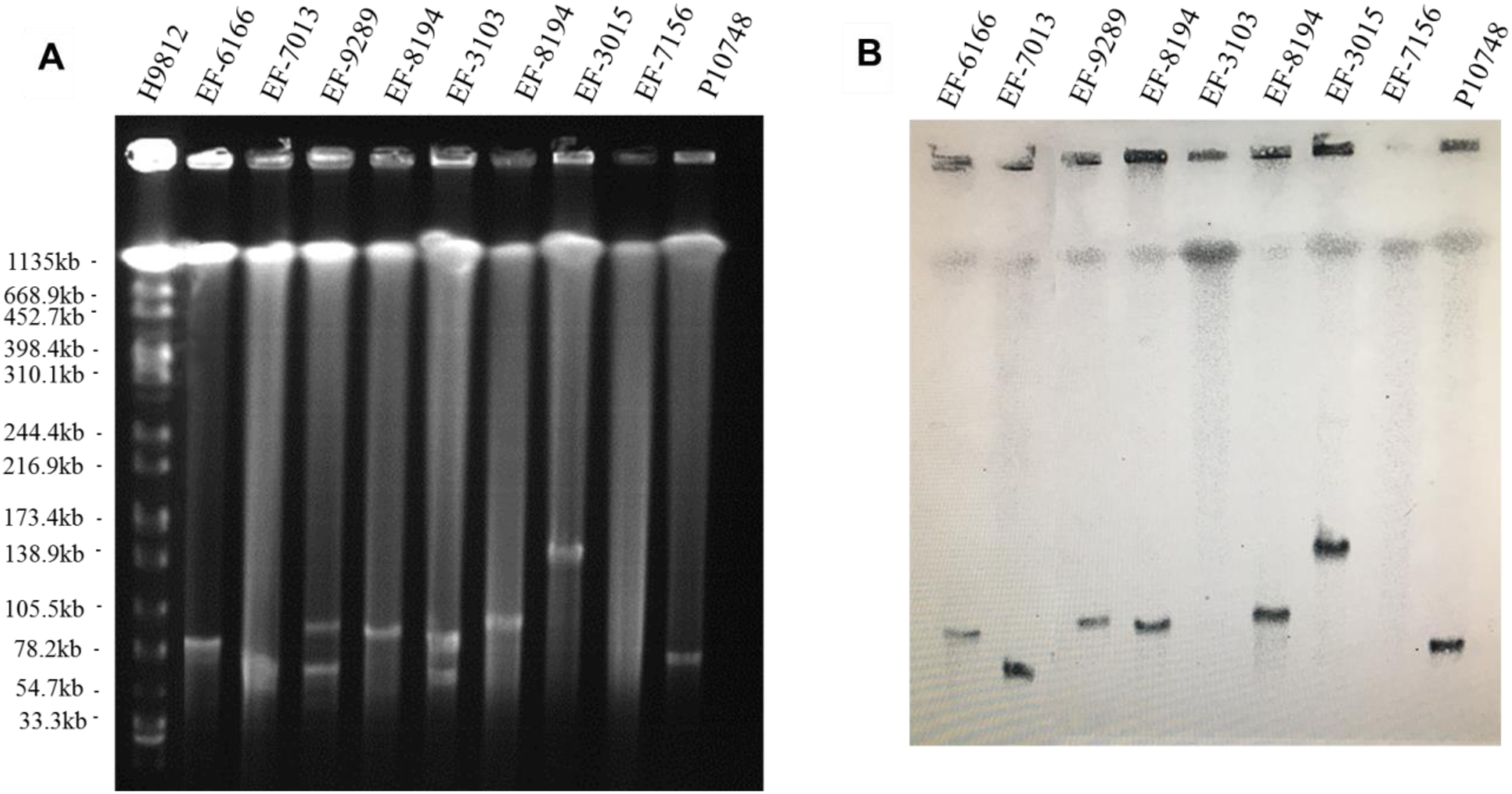
Determination of the location of *optrA* in linezolid-resistant *E. faecalis* isolates. (**A**) Representative results of S1 nuclease-pulsed-field gel electrophoresis analysis. The first lane contained *XbaI*-digested chromosome of *Salmonella* H9812 as a DNA size marker. (**B**) Representative results of Southern hybridization showing the location of *optrA* in plasmids or chromosomes. Individual isolates analyzed are indicated at the top.

### 3.2. Horizontal transmission of *optrA* gene

In conjugation experiment with *E. faecalis* JH2-2 as a recipient, positive transfer of *optrA* gene was achieved in 23/26 (88.5%) *E. faecalis* clinical isolates with *optrA*-carrying plasmids. The other 3 isolates showed no transfer despite multiple attempts. None of the 18 isolates containing *optrA* in the chromosome showed positive transfer. In drug susceptibility testing of the 23 transconjugants, the MIC value of linezolid changed from susceptibility (2 mg/L) to resistance (8 mg/L) and *E. faecalis* ATCC 29212 was used for the control strain. In addition, these 23 transconjugants exhibited resistance to chloramphenicol, clindamycin, erythromycin, and tetracycline compared with the recipient strain *E. faecalis* JH2-2 (data not shown). By PCR, in each of the 23 transconjugants we detected multiple resistance genes, including *optrA* (23/23, 100%), *tetL* (23/23, 100%), *tetM* (11/23, 47.8%), *ermB* (22/23, 95.6%), *lnuB* (20/23, 87.0%) and *fexA* (22/23, 95.6%) (Table1).

### 3.3. Detection of *rep*-families in *E. faecalis* transconjugants and their original clinical isolates

Four types of plasmids were identified by PCR in 20 transconjugant isolates carrying *optrA* while other three isolates did not show any of the 19 plasmid types targeted in our study (Table 1). The dominant plasmid type was *rep*_9_ family (prototype pCF10), which was detected in 78% (18/23) of isolates. *Rep*_1_ (prototype pIP501) accounted for the second major *rep*-family being identified in 39% (9/23) isolates. *rep*_7_ (pIP501 prototype) and *rep*_6_ (pIP501 prototype) were detected with lower frequencies (17.4% or 4/23 and 8.7% or 2/23, respectively). Other plasmids including *re_p_*_2-5_ *rep*_8_, *rep*_10-11_, *rep*_13-16_, *rep*_18-19_ and the *rep_Unique_* were not detected in any isolates.

Of the 20 successfully typed isolates, 9 contained a single plasmid type, 9 contained a mixture of two plasmid types and the remaining 2 contained a mixture of three plasmid types (Table 1).

Of the 23 original clinical isolates showing positive transfer of *optrA*-carrying plasmids, 18 contained a *rep_9_* type of plasmids as determined by PCR using primers specific for *rep_9_*-type plasmid.

### 3.4 Characteristics of the *optrA-carrying* plasmid in *E. faecalis* strain P10748

*De novo* assembly of Illumina HiSeq data from *E. faecalis* P10748 resulted in a complete genome of plasmid pEF10748 containing 53,178 bp (GenBank accession no. KPMK993385). This size is consistent with the results of Southern blot (Fig. 1), supporting the correct assembly of the plasmid genome. This genome has a GC content of 35%, similar to that of the *E. faecalis* genome. A total of 58 coding sequences (CDSs) were identified, with majority of them encoded in the same orientation, similar to known pheromone plasmids in *E. faecalis,* such as pCF10 (43) and pAD1 (44). The predicted genes of all CDSs are listed in Table 2 and Fig. 2.

**Table 2.**
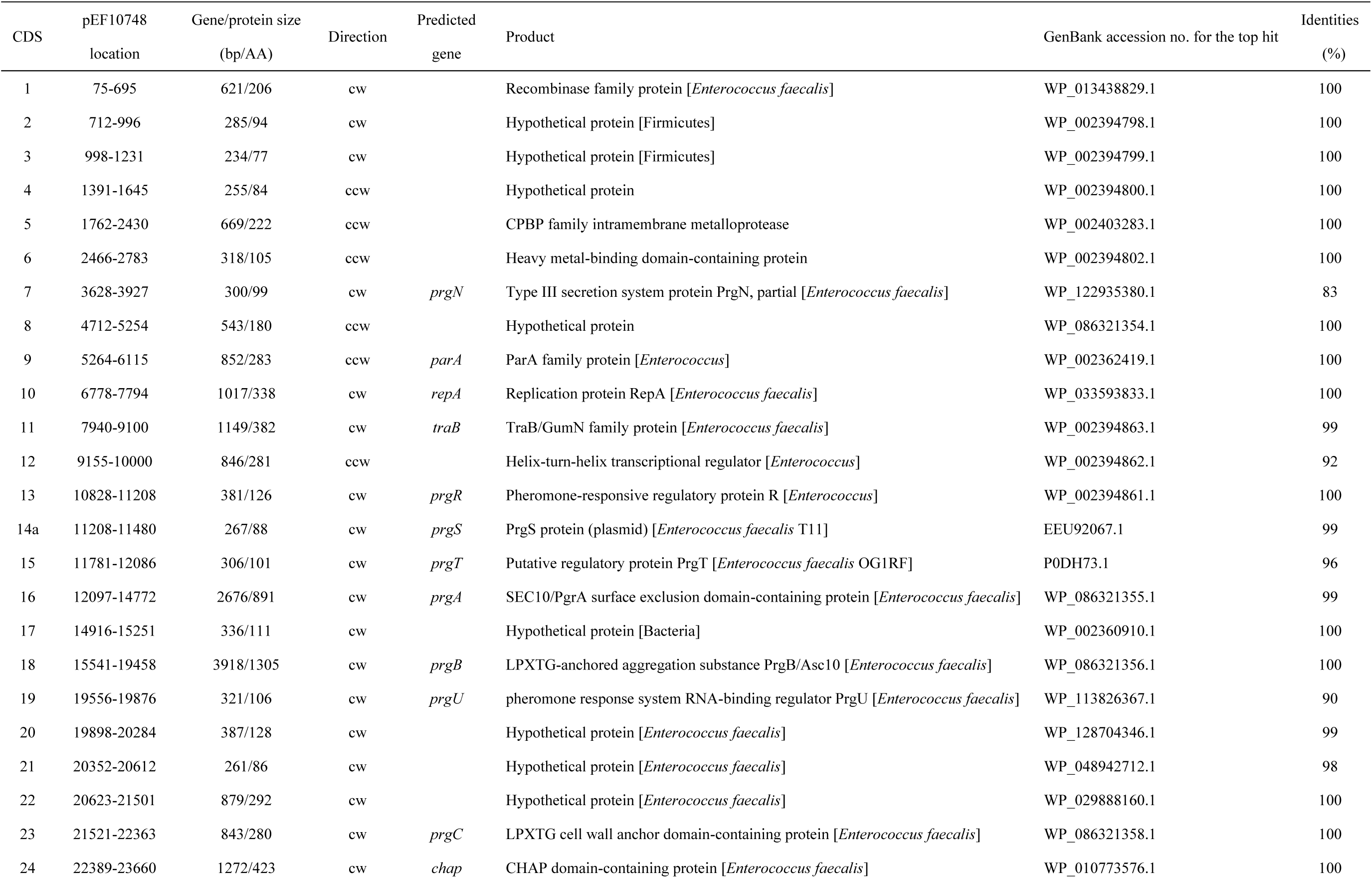
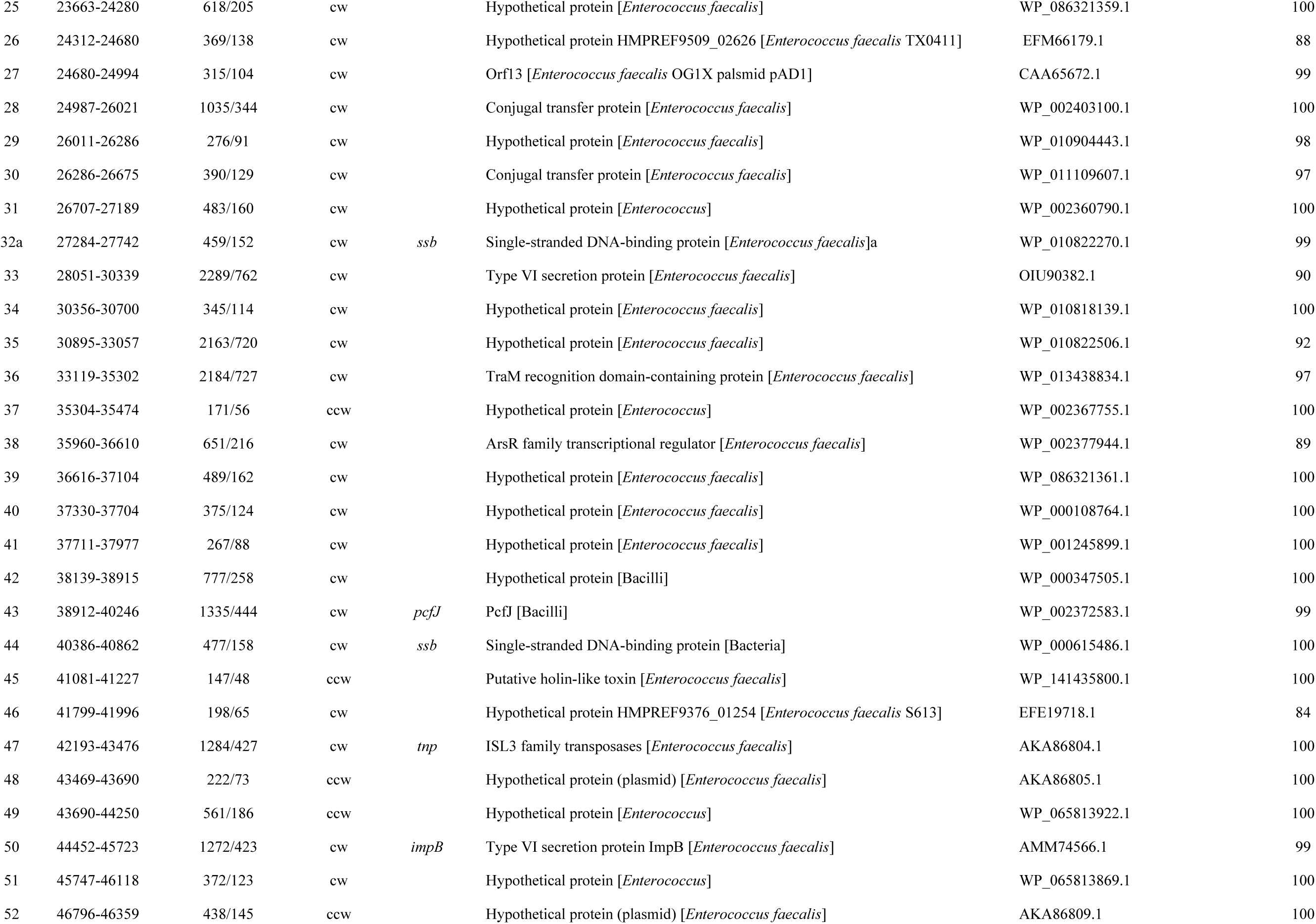
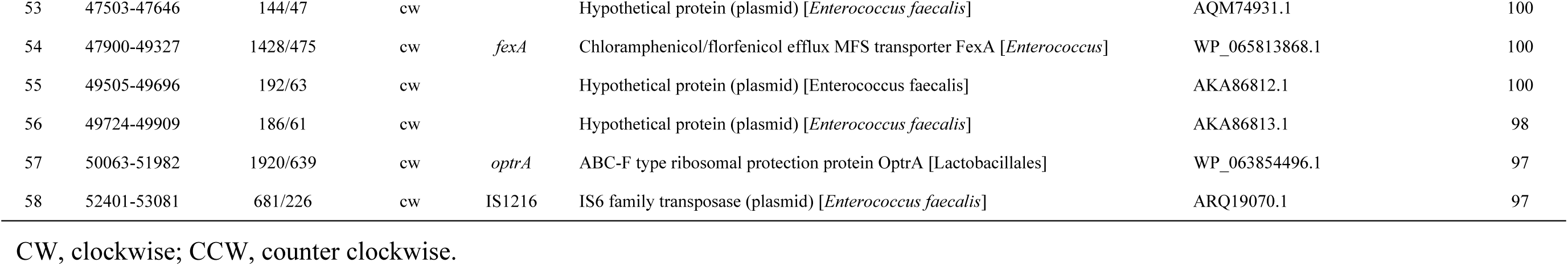
Genes encoded on plasmid pEF10748.

**Figure 2.**
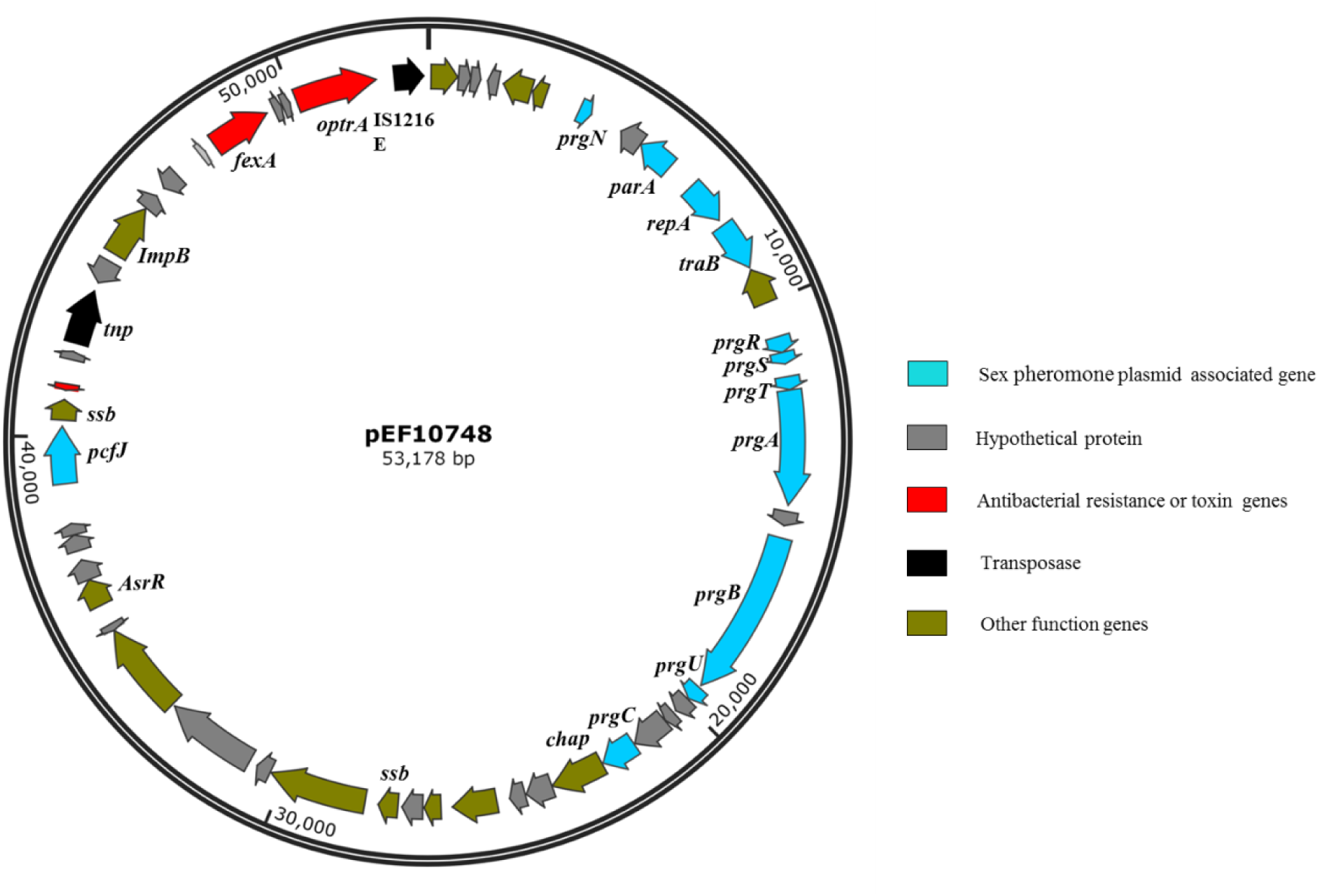
Gene organization in plasmid pEF10748 from the *E. faecalis* clinical isolate P10748. Arrows indicate the CDSs and their transcription directions. The putative functions of CDSs are color-coded as indicated at the right.

The full-length sequence of plasmid pEF10748 was not identical to any plasmid sequences currently available in GenBank while being most closely related to plasmid pKUB3006-1 in an *E. faecalis* isolate reported from Japan (45), with 78% coverage and 99% sequence identity for a ∼42 kb region (Table S3). Other plasmids with significant overlap to this region included pTEF1, pAD1, pMG2200, pCF10 and pTEF2 (Fig. 3). All these plasmids were identified in *E. faecalis* from human and swine samples and contain multiple sex pheromone response genes, including *traB*, *prgA*(*sea1*), *prgB* (*asa1*), *prgC, prgU*, *prgR*, *prgS* and *prgT*.

**Figure 3.**
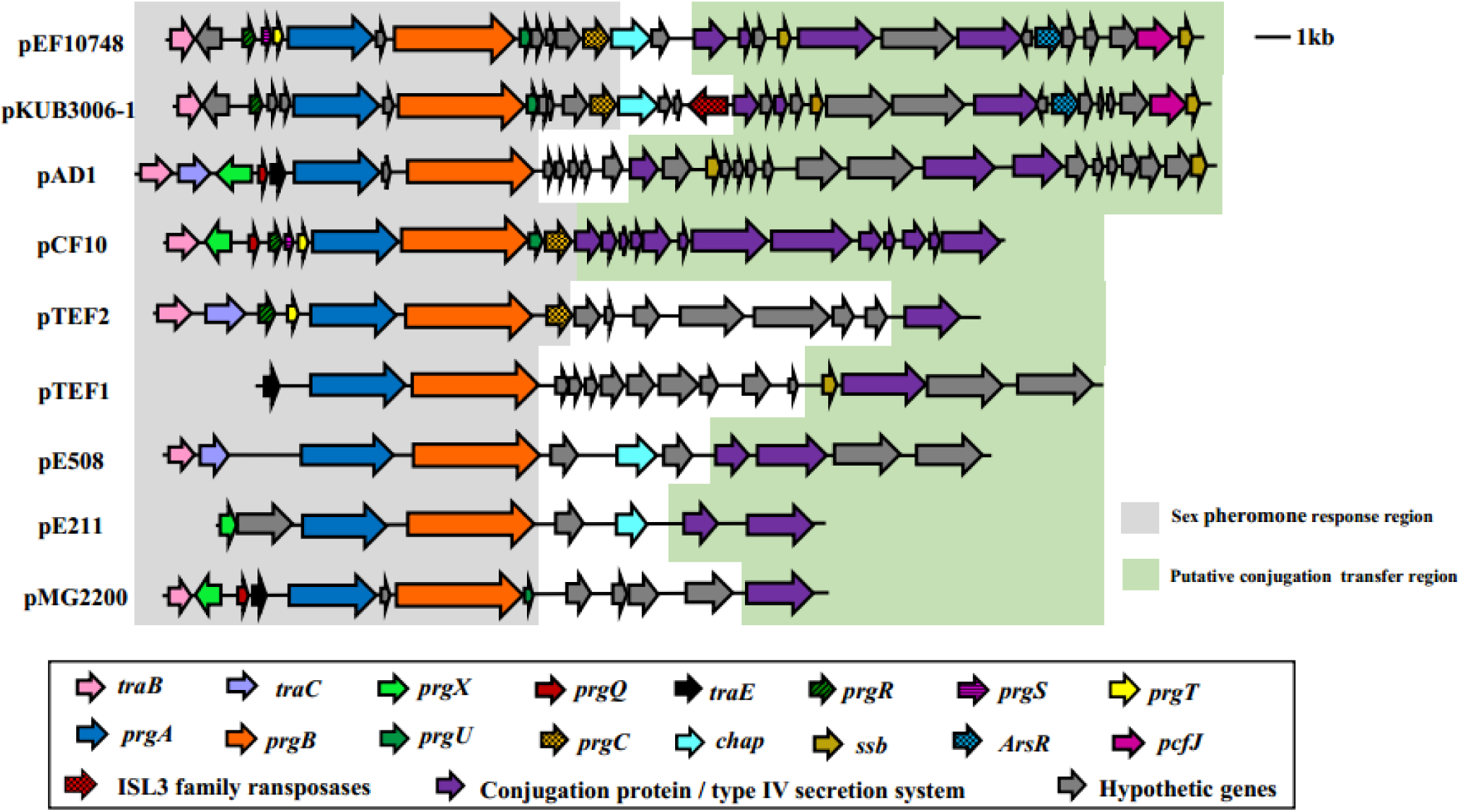
Comparison of the sex pheromone gene cluster among known sex pheromone plasmids. Different genes are color-coded as shown in the box on the bottom. Of note, for the region shown, the gene organization in plasmid pEF10748 identified in this study is most similar to that of pKUB3006-1 reported by Kuroda *et al.* (45). The bacterial origin and GenBank accession numbers of all plasmids shown are available from Supplementary Table S3.

The *optrA* gene in plasmid pEF10748 was surrounded by chloramphenicol/florfenicol resistance gene *fexA,* ISL3 transposase, *Imp* and IS1216E element, all of which were located in a ∼11 kb region (Table 2 and Fig. 2). This region showed highest homology (>99.9% sequence identity) to the following plasmids: pXY17, p10-2-2, p29462, p1202 and pE121 (Table S3 and Fig. 4). All these plasmids were identified from *E. faecalis* in China.

**Figure 4.**
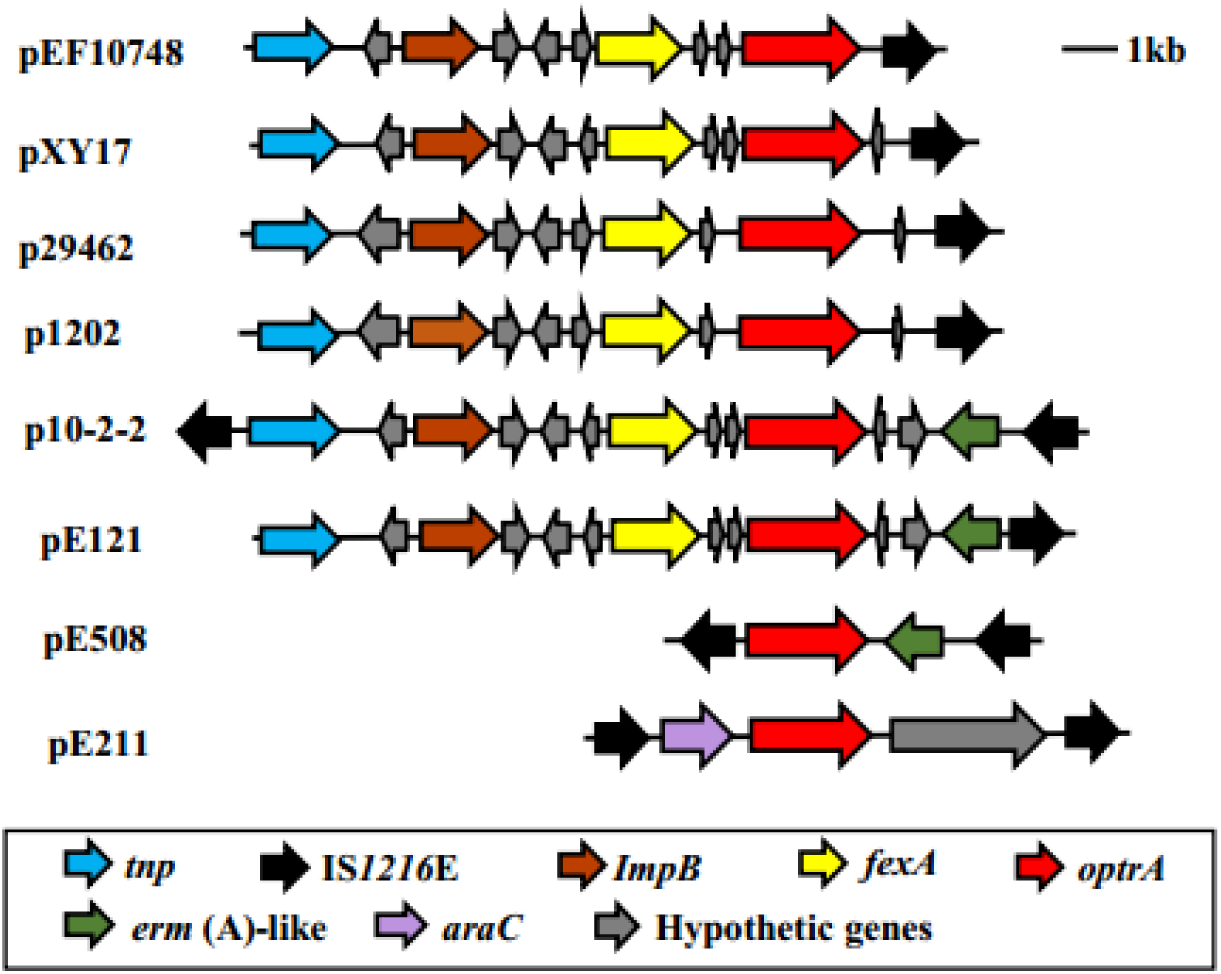
Comparison of the genetic environment of the linezolid resistance *optrA* among different plasmids. Different genes are color-coded as shown in the box on the bottom. Of note, for the region shown, the gene organization in plasmid pEF10748 identified in this study is most similar to that of pXY17 reported by He *et al*. (38). The bacterial origin and GenBank accession numbers of all plasmids shown are available from Supplementary Table S3.

### 3.5. Distribution of key sex-pheromone-response genes in *optrA*-carrying *E. faecalis* isolates and their transconjugants

Among the 23 *E. faecalis* isolates showing a positive transfer of *optrA*-carrying plasmids, 15 were PCR-positive for the *prgA* gene with both the original clinical isolates and their respective transconjugants, 5 were positive with the original isolate only, and the remaining 3 were negative with both the original isolates and transconjugants (Table 3). In PCR testing for the *prgB* gene, 21 were positive with both the original isolates and transconjugants, and the remaining 2 were positive with the original isolate only. In PCR testing for the *prgC* gene, 10 were positive with both the original isolates and transconjugants, 5 were positive with the original isolate only, and the remaining 8 were negative with both the original isolates and transconjugants. Overall, all 23 isolates were positive for at least one of the three key pheromone-response genes with either the original isolates or transconjugants; only 7 isolates were positive for all 3 genes with both the original isolates and transconjugants. *prgB* was the most common gene, which was detected in all 23 original isolates.

**Table 3.** Distribution of key sex-pheromone-response genes and results of clumping assay in 23 *E. faecalis* isolates showing successful transfer of *optrA*-carrying plasmids.

|  | *rep*-type | Pheromone-response genes<sup>a</sup> |  |  |  | Clumping assay<sup>b</sup> |  |
| --- | --- | --- | --- | --- | --- | --- | --- |
|  |  | *rep9* | *prgA* | *prgB* | *prgC* | cCF10 | cAD1 |
| p10748 | 9 | +/+ | +/+ | +/+ | +/+ | ++ | + |
| EF-3015 | 6, 9 | +/+ | -/+ | +/+ | -/- | + | ++ |
| EF-7013 | 9 | +/+ | -/+ | +/+ | +/+ | +++ | +++ |
| EF-8194 | 9 | +/+ | +/+ | +/+ | +/+ | +++ | +++ |
| EF-9289 | 1, 7, 9 | +/+ | +/+ | +/+ | -/- | ++ | +++ |
| EF-3139 | 1, 9 | +/+ | +/+ | +/+ | +/+ | - | + |
| EF-2084 | 9 | +/+ | +/+ | +/+ | -/+ | - | - |
| EF-0132 | 1, 9 | +/+ | -/+ | +/+ | -/- | - | - |
| EF-4003 | 1, 9 | +/+ | +/+ | +/+ | +/+ | + | - |
| EF-8042 | 7 | -/- | +/+ | +/+ | -/- | - | - |
| EF-0074 | 1, 9 | +/+ | +/+ | +/+ | +/+ | - | - |
| EF-6165 | 7, 9 | +/+ | +/+ | +/+ | +/+ | +++ | +++ |
| EF-7094 | 1 | -/- | +/+ | +/+ | +/+ | - | - |
| EF-1127 | 9 | +/+ | -/+ | +/+ | +/+ | - | - |
| EF-8137 | 1, 9 | +/+ | -/- | +/+ | -/- | + | - |
| EF-8282 | 1, 7, 9 | +/+ | +/+ | +/+ | -/+ | + | + |
| EF-4245 | 6,9 | +/+ | +/+ | +/+ | -/+ | + | - |
| EF-1080 | 9 | +/+ | -/- | +/+ | -/+ | - | - |
| EF-2216 | 9 | +/+ | -/- | +/+ | -/- | + | - |
| EF-6166 | 1,9 | +/+ | -/+ | -/+ | -/+ | - | - |
| EF-5136 | NA<sup>c</sup> | -/- | +/+ | +/+ | -/- | - | - |
| EF-5015 | NA<sup>c</sup> | -/- | +/+ | -/+ | +/+ | - | - |
| EF-2021 | NA<sup>c</sup> | -/- | +/+ | +/+ | -/- | - | - |

<sup>a</sup>The presence (+) or absence (-) in transconjugants/original clinical isolates by PCR.
<sup>b</sup>Results of clumping (with original isolates only) were scored as: -, no clumping; +, weak clumping; ++, moderate clumping; +++, strong clumping.
NA, not available in plasmid typing with 19 known *rep* families (Table 1).

Of the 18 original isolates containing a *rep_9_* type of plasmids, all were positive for *prgB*, 15 were positive for *prgA* and 13 were positive for *prgC*. On the other hand, of the 5 original isolates that did not carry a *rep_9_* type of plasmids, and all were positive for *prgA* and *prgB* while only 2 were positive for *prgC*.

### 3.6. Determination of sex pheromone responses by CIA

In CIA assay with synthetic pheromones cAD1 and cCF10 as inducers, 12 of 23 *optrA*-carrying isolates showed different degrees of clumping (Fig. 5) with at least one inducer, including 3 showing very strong clumping and 9 showing weak to moderate clumping (Table 3). None of the 5 isolates without a *rep_9_*-type plasmid showed apparent clumping. In CIA assay with JH2-2 filtrates, 17 of 23 *optrA*-carrying isolates (all containing *rep_9_*-type plasmids) displayed apparent aggregation under microscopy while the remaining 6 isolates showed no aggregation.

**Figure 5.**
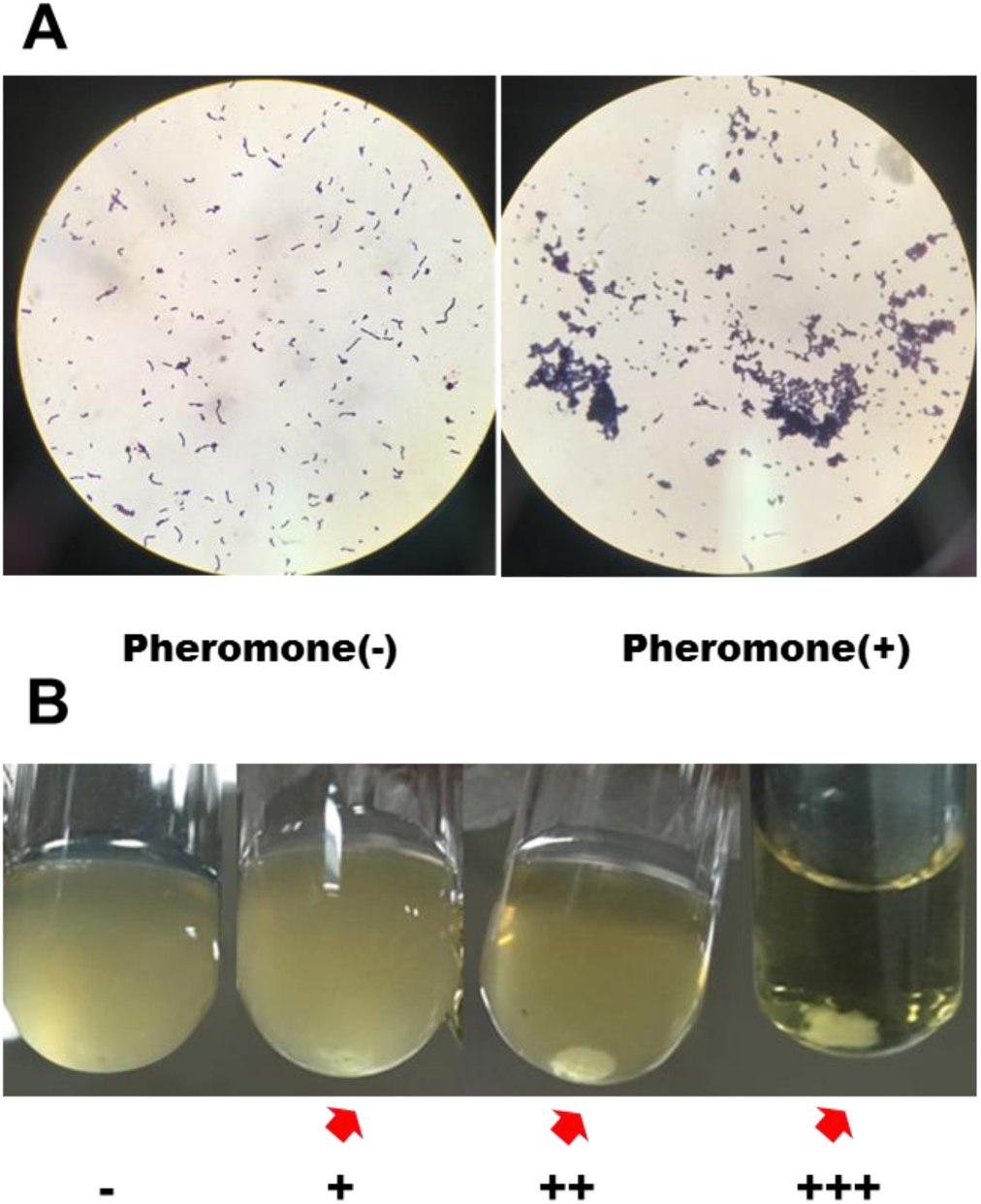
Representative results of clumping assay induced by sex pheromones. (**A**) Microscopic view of *E. faecalis* strain P10748 mounted on glass slide after growing for 3h in the absence (-) or presence (+) of sex pheromones produced by *E. faecalis* JH2-2. Magnification at X1,000. (**B**) *E. faecalis* strain EF-2084, EF-3015, P10748, EF-6165 grown in culture tubes in the presence of the synthetic pheromone cCF10. Different degrees of cell clumping were indicated by red arrows: negative (-), weak (+), moderate (++) and strong (+++).

## DISCUSSION

In this study, we performed comprehensive molecular and phenotypic profiling of 44 *E. faecalis* clinical isolates with linezolid-resistance through combination of PFGE, DNA hybridization, PCR-based genotyping, whole plasmid genome sequencing, antibiotic susceptibility test, bacterial conjugation and clumping induction assay. Our results strongly suggest that *optrA*-mediated linezolid-resistance can be widely disseminated through sex pheromone plasmid transfer.

Our conclusion on the sex pheromone plasmid transfer mechanism is supported by the following findings. First, we detected a high prevalence of plasmid-borne *optrA* gene in *E. faecalis* clinical isolates. The proportion of isolates carrying *optrA* in the plasmid was slightly higher isolates carrying *optrA* in the chromosome (59% *vs* 41%), which is consistent with previous reports in China (13, 15) and other countries (6, 22, 37). Location of the resistance gene is a plasmid is known to enable fast and efficient gene transfer within and between different bacterial host species. Second, we detected a high transferability (23/26 or 88%) of *optrA*-carrying plasmids based on studying a case series of 26 *E. faecalis* clinical isolates with linezolid resistance and carrying *optrA* in plasmids (Table 1). This transferability was confirmed by PCR and MIC testing of all transconjugants. To our knowledge, this study is the largest to date to assess the transferability of *optrA*-carrying plasmids in clinical isolates of *E. faecalis.* Third, sequencing of the full plasmid genome in one *E. faecalis* clinical strain (P10748) confirmed the co-localization of *optrA* with almost all known sex-pheromone response genes, including the typical *prgA-prgB-prgC* cassette, in the same plasmid (Fig. 2). Plasmids with such gene organization have not been reported previously from any *optrA*-carrying *E. faecalis* isolates. While there has been one report of two pheromone-responsive plasmids carrying *optrA* in *E. faecalis* (30), both plasmids were identified in *E. faecalis* isolates from pigs, with only one of them (pE508) containing a single sex pheromone gene (*prgB*) based on full plasmid genome sequencing. Fourth, our plasmid typing studies demonstrated, for the first time, the abundant presence of *rep*_9_-type plasmids (prototype pCF10) in *optrA*-carrying *E. faecalis* clinical isolates (17/26 or 65%, Table 1), implying a high prevalence of sex pheromone plasmids in clinical isolates. Consistent with this observation, our PCR-based genotyping revealed the presence of three key sex pheromone response genes (*prgA, prgB* and *prgC*) in almost all *optrA*-carrying *E. faecalis* clinical isolates (Table 3). Fifth, our functional studies with CIA detected different degrees of cell clumping in most of the 23 *E. faecalis* clinical isolates carrying *optrA* in plasmids, indicating that pheromone-inducible conjugation is operational in these isolates. The absence of clumping in some of these isolates may be caused by the presence of other pheromone receptors different from those for cAD1 and cCF10 or JH2-2 infiltrate used in the CIA assay. Finally, the hypothesis of sex pheromone-mediated transfer of *optrA* is further supported by our previous transcriptomics (31) and proteomics (32) studies of the *E. faecalis* P10748, which consistently showed that OptrA and several sex pheromone response molecules (PrgA, PrgB and PrgY) were among the most significantly up-regulated molecules.

The plasmid pEF10748 identified in this study represents a novel plasmid for *E. faecalis*. In addition to the presence of *OptrA* and multiple sex pheromone response genes described above, this plasmid contains other drug resistance and virulence determinants. The *fexA* gene, which confers resistance to chloramphenicol and florfenicol, is located closely to the *optrA* gene in pEF10748. The region containing these two drug resistance genes is flanked by two transposase genes including the IS1216 and ISL3 family transposases (Fig. 2). The same gene organization has been reported in only one partially sequenced plasmid (pXY17) from an *E. faecalis* clinical isolate (38). It is likely that this organization will facilitate the movement of the resistance genes to different locations (plasmids or chromosomes) or different bacteria. This possibility awaits further investigation. In addition to drug resistance genes, plasmid pEF10748 encodes a putative holin-like toxin, which has been previously identified in seven plasmids from different *E. faecalis* strains based on GenBank search. As holin proteins are known to function through disruption of the host cell membrane (46), we speculate that this protein may play a role in gene transfer by releasing mobile genetic elements. Co-localization of these resistance and virulence determinants with a set of multiple sex pheromone response genes may enable a highly efficient dissemination of these mobile genetic materials.

This study clearly has some limitations. The distribution of key sex pheromone response genes in clinical isolates was determined by PCR alone using total genomic DNA, which is unable to distinguish whether these genes are located in plasmids. Clarification of this question requires further studies by DNA hybridization or full plasmid genome sequencing. Despite the availability of a relative large collection of *optrA*-carrying plasmids, we sequenced only one of them (strain P10748), which was chosen due to our previous studies by transcriptome and proteome analysis (31, 32). It is unknown if the remaining plasmids also carry sex pheromone response genes as well as other mobile genetic elements. We anticipate that continuing decrease in NGS cost should allow us to sequence all these plasmids as well as related chromosomes.

## Conclusion

This report represents the largest study of the prevalence and transferability of *optrA*-carrying plasmids in *E. faecalis* clinical isolates, and the first identification of a plasmid carrying *optrA* along with multiple sex pheromone response genes in clinical isolates. Our integrative molecular and phenotypic analysis strongly suggests that *optrA*-mediated linezolid-resistance can be widely disseminated through sex pheromone plasmid transfer. Further studies are needed to determine how this transfer is regulated and what is the best strategy to monitor and control the transmission.

## Acknowledgments

We would like to thank Xiuli He and Yuhang Liu for their assistance in collection of clinical samples. We also wish to thank Dr. Liang Ma at the National Institutes of Health, Bethesda, Maryland for critical reviewing of the manuscript.

## Declarations

### Ethical Approval

For this type of study (with no human subjects involved), formal consent is not required.

---

**Supplementary Table S1.** Resistance phenotypes of 26 clinical *E. faecalis* isolates carrying *optrA* used in this study.

| Isolate ID | Resistance phenotypes<sup>a</sup> |
| --- | --- |
| P10748 | CIP, MOX, LEV, TET, ERY, GEN, CLI,STR, LZD, DAF |
| EF-3015 | TET, ERY, GEN, CLI, STR, LZD, DAF |
| EF-7013 | CIP, MOX, LEV, TET, ERY, CLI, GEN, STR, LZD, DAF |
| EF-8194 | CIP, MOX, LEV, TET, ERY, GEN, CLI, STR, LZD, DAF |
| EF-9289 | PEN, TET, ERY, GEN, CLI, LZD, DAF |
| EF-3139 | PEN, TET, ERY, STR, GEN, CLI, LZD, DAF |
| EF-5136 | TET, ERY, CLI, LZD, DAF |
| EF-2084 | CIP, MOX, LEV, TET, ERY, GEN, CLI, STR, LZD, DAF |
| EF-0132 | CIP, MOX, LEV, TET, ERY, GEN, CLI, STR, LZD, DAF |
| EF-4003 | CIP,MOX,LEV,TET,ERY,GEN,CLI,STR,LZD,DAF |
| EF-8042 | CIP, MOX, LEV, TET, ERY, CLI, LZD, DAF |
| EF-0074 | CIP, MOX, LEV, TET, ERY, GEN, CLI, STR, LZD, DAF |
| EF-6165 | PEN, CIP, LEV, TET, ERY, STR, CLI, LZD, DAF, FUR |
| EF-7094 | TET, ERY, GEN, CLI, LZD, DAF |
| EF-1127 | TET, ERY, STR, GEN, CLI, LZD, DAF |
| EF-8137 | CIP, MOX, LEV, TET, ERY, GEN, CLI, LZD, DAF |
| EF-8282 | CIP, TET, ERY, STR, CLI, LZD, DAF |
| EF-4245 | CIP, LEV, TET, ERY,STR, CLI, LZD, DAF |
| EF-1080 | CIP, MOX, LEV, TET, ERY, CLI, STR, LZD, DAF |
| EF-2216 | TET, ERY, CLI, LZD, DAF |
| EF-6166 | CIP, MOX, LEV, TET, ERY, GEN, CLI, LZD, DAF |
| EF-5015 | CIP, MOX, LEV, TET, ERY, GEN, CLI, STR, LZD, DAF |
| EF-2021 | CIP, MOX, LEV, TET, ERY, CLI, GEN, LZD, DAF |
| EF-8014 | CIP, MOX, LEV, TET, ERY, GEN, CLI, STR, LZD, DAF |
| EF-3186 | TET, ERY, CLI, LZD, DAF |
| EF-1090 | CIP, MOX, LEV, TET, ERY, GEN, CLI, STR, LZD, DAF |

<sup>a</sup>Determined in our previous study (33).
AMP, ampicillin; PEN, penicillin G; CLI, ciprofloxacin; MOX moxifloxacin; LEV, levofloxacin; TET, tetracycline; ERY, erythromycin; STR, high-level streptomycin resistance; GEN, high-level gentamicin resistance to chemotherapy; CLI, clindamycin; DAF, quinupristin/dalofopine; FUR, nitrofurantoin; LZD, linezolid.

**Supplementary Table S2.**
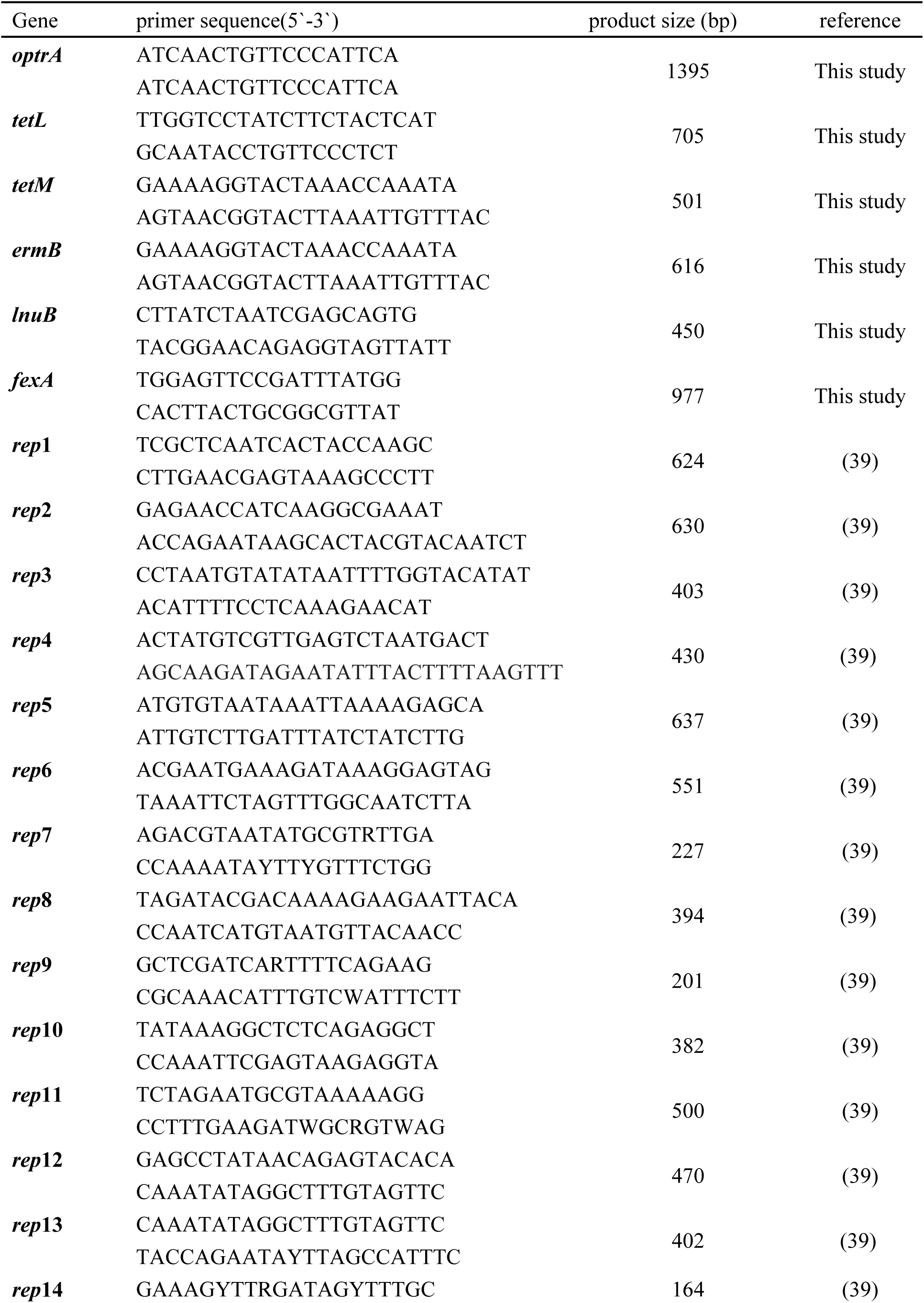
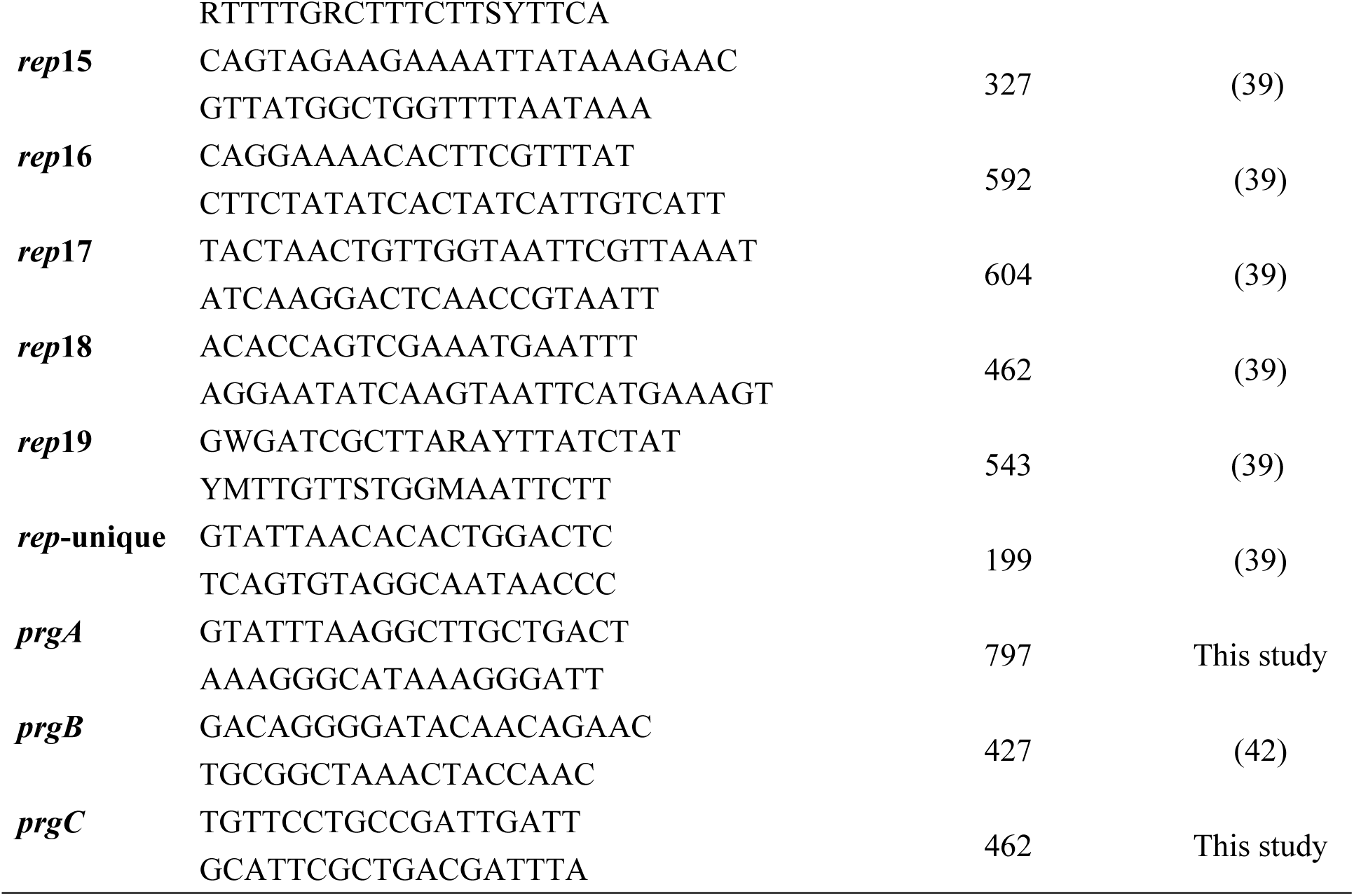
Primers used for PCR

**Supplementary Table S3.**
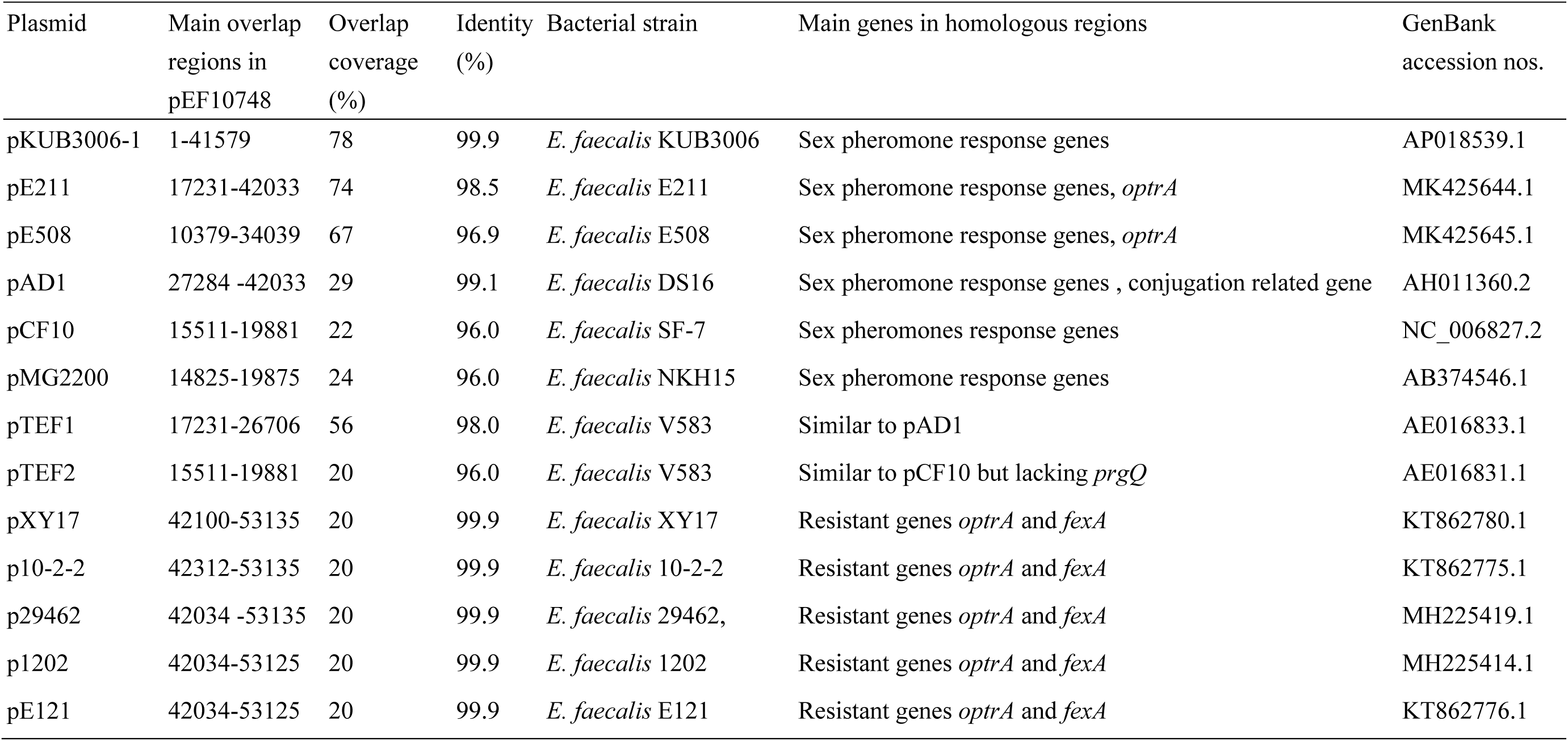
Homology of plasmid pEF10748 with selected known plasmids in *E. faecalis*.

